# A study of parietal-motor connectivity by intraoperative dual cortical stimulation

**DOI:** 10.1101/747337

**Authors:** Luigi Cattaneo, Davide Giampiccolo, Pietro Meneghelli, Vincenzo Tramontano, Francesco Sala

**Author notes:** **Corresponding author:** Luigi Cattaneo, MD, PhD, Institute of Neurophysiology, Dept. of Neuroscience, Biomedicine and Movement, University of Verona, Piazzale L.A. Scuro 10, 37134 Verona, Italy.

## Abstract

the function of the primate’s posterior parietal cortex in sensorimotor transformations is well-established, though in humans its complexity is still challenging. Well-established models indicate that the posterior parietal cortex influences motor output indirectly, by means of connections to the premotor cortex, which in turn is directly connected to the motor cortex. The possibility that the posterior parietal cortex could be at the origin of direct afferents to M1 has been suggested in humans but has never been confirmed directly. In the present work we assessed during intraoperative monitoring of the corticospinal tract in brain tumour patients the existence of short-latency effects of parietal stimulation on corticospinal excitability to the upper limb. We identified several foci within the inferior parietal lobule that drove short-latency influences on cortical motor output. Active foci were distributed along the postcentral gyrus and clustered around the anterior intraparietal area and around the parietal operculum. For the first time in humans, the present data show direct evidence in favour of a distributed system of connections from the posterior parietal cortex to the ipsilateral primary motor cortex.

## Introduction

### The role of the posterior parietal cortex in active behaviour

The last 40 years have witnessed a radical change in our view of the parietal cortex (Mountcastle *et al*., 1975). The posterior parietal cortex, once labelled as “associative cortex” is now well-known for receiving multimodal sensory information and integrating it into a praxic, behaviourally-committed representation of the world around us. Solid evidence in the field of neuropsychology, neuroimaging and neurostimulation indicates that the posterior parietal cortex is necessary for goal-directed behaviour. Symptoms frequently caused by lesions of the parietal lobe include deficits in sensorimotor processes, such as optic ataxia (Andersen *et al*., 2014) or apraxia (Goldenberg, 2009). Direct stimulation of the human parietal cortex has been shown to produce movements in all body segments (Penfield and Boldrey, 1937; Balestrini *et al*., 2015). Current evidence indicates in the human superior parietal lobule the machinery for sensorimotor transformation in spatially-oriented movements (for reviews see Culham and Valyear, 2006, Filimon, 2010 and Gallivan and Culham, 2015) controlling also some aspects of distal prehension movements (Monaco *et al*., 2015; Cavina-Pratesi *et al*., 2018). Visual features of objects, used to guide distal, object-directed movements are represented in humans in the anterior intraparietal region (Culham *et al*., 2003; Frey *et al*., 2005; Begliomini *et al*., 2007; Grol *et al*., 2007; Stark and Zohary, 2008; Hinkley *et al*., 2009; Verhagen *et al*., 2012; Orban, 2016). The role of the inferior parietal lobule in movement is less clear. Grasping-related activity in the anterior intraparietal region is found along the descending part of the precentral sulcus, up to the parietal operculum with some specialization for tool use of the more ventral areas (Orban, 2016). More ventrally, the parietal opercular region is also thought to be a site of sensorimotor integration, mainly in the somatosensory modality (Eickhoff *et al*., 2006b, *a*, 2010). Summing up, imaging data in humans indicate an extended region ranging from the superior parietal lobule to the parietal operculum involved in sensorimotor processes and specialized in distinct functional aspects.

### Parietal-motor pathways

How does the motor cortex use motor-relevant information from the posterior parietal cortex? The influence of the parietal cortex on motor output is generally considered to be indirect. According to influential models, based on monkey anatomy, the parietal cortex modulates corticospinal activity in an indirect way, through the premotor cortex (Murata *et al*., 1997; Wise *et al*., 2002; Rizzolatti *et al*., 2014; Kaas and Stepniewska, 2016). However, even anatomical data in monkeys are controversial in this respect. The existence of direct, monosynaptic connections from the posterior parietal cortex to the upper limb representation of the primary motor cortex has been demonstrated by several independent works (Strick and Kim, 1978; Rozzi *et al*., 2006; Bruni *et al*., 2018). Such data offer the anatomical bases for a possible direct pathway by which the posterior parietal cortex might control directly corticospinal output. In addition to this, it has been recently shown that the posterior parietal cortex of macaques has a direct access to spinal motor neurons by means of corticospinal axons (Rathelot *et al*., 2017). In humans, several recent lines of evidence have suggested that the posterior parietal cortex might have a more direct influence on motor output. Non-invasive brain stimulation, i.e. transcranial magnetic stimulation (TMS) suggests that the parietal cortex could give origin to direct cortico-cortical connections to the primary motor cortex (M1) (Koch *et al*., 2007, 2008b, 2010; Ziluk *et al*., 2010; Cattaneo and Barchiesi, 2011; Karabanov *et al*., 2013; Maule *et al*., 2015), involved in skilled upper limb movements. Summing up, there is ample evidence in both nonhuman and human primates to support the possibility that the parietal lobe could modulate corticospinal output also directly through M1, besides the well-established indirect pathway through a relay in the premotor cortex (see Koch and Rothwell (2009) and Vesia and Davare (2011) for a review of parieto-M1 interaction models). The active role of the posterior parietal cortex in producing movements is being currently re-evaluated as a potential source of “pre-motor” afferents to the premotor cortex, where pre-motor is used here in a functional sense rather than anatomical. Imaging data provided indirect evidence of posterior parietal-motor anatomical and functional connectivity (Guye *et al*., 2003; Koch *et al*., 2010; Yin *et al*., 2012). However, direct evidence in favour of direct connections between the parietal and the motor cortex in humans are lacking.

### Testing direct parieto-motor pathways intraoperatively

We aim to fill this gap in current knowledge with the present work in which we tested cortico-cortical connectivity by means of intra-operative direct cortical stimulation (DCS) with a dual-pulse paradigm similar to that employed with dual-coil TMS (Koch and Rothwell, 2009). Supra-threshold test stimuli were delivered to M1 and the resulting motor evoked potentials (MEPs) were systematically recorded form distal upper limb muscles, and in some cases from facial and lower limb muscles. In some trials a conditioning stimulus was delivered to different regions of the parietal cortex at variable inter-stimulus intervals (ISIs) ranging from 4ms to 16 ms. In some other trials, only conditioning stimuli were delivered. The conditioning stimulus itself does not activate the corticospinal motor pathways, as witnessed by the systematic absence of MEPs in such trials. The modulation of motor output by conditioning stimuli is generally considered as evidence of cortico-cortical functional connectivity between the target of conditioning stimuli and the motor cortex. It is important to note that a necessary pre-requisite for the realization of dual-stimulation paradigms is that the corticospinal tract must be activated trans-synaptically by the test stimuli, because direct axonal stimulation of corticospinal axons produces MEPs that arise downstream of the putative site of interaction between the conditioning and the test stimuli. In this respect, the information currently available indicates that direct cortical stimulation (DCS) is effective also under general anaesthesia in exciting cortical output trans-synaptically, also at low stimulation intensities (Katayama *et al*., 1988; Hanajima *et al*., 2002; Yamamoto *et al*., 2004; Lefaucheur *et al*., 2010). Invasive intra-operative monitoring (IONM) in neurosurgery therefore offers the unique opportunity to assess cortico-cortical connectivity *in vivo*, with extraordinary spatial resolution and anatomical precision. Indeed, the results of the present work indicated a diffuse field of parietal spots exerting short-latency modulation of corticospinal output in the range of 6-15 ms of inter-stimulus intervals, distributed along the post-central sulcus. Such active spots showed higher density in two large clusters in the anterior intraparietal region and in the parietal operculum. We show that the anterior portion of the inferior parietal lobule and intraparietal region can be functionally considered as a “pre-motor” region. The behavioural significance of such connections is yet to be determined.

## Methods

### Patients

The study proposal is in accordance with ethical standards of the Declaration of Helsinki. All stimulations and recordings were performed in the context of clinical intraoperative neurophysiological monitoring (IONM). Patients scheduled for tumour removal in the vicinity of the parietal cortex were screened for enrolment. The inclusion criteria were: (1) brain tumour necessitating intraoperative neurophysiological monitoring (2) over 18 years of age. Exclusion criteria were (1) extended cortico-subcortical damage to the parietal lobe (2) voluntary decision of the patient not to be included in the cohort. Seventeen patients (age 39-79; 10M-7F; 17 righthanded) were included in this study, recruited from the Verona University Hospital. Patient’s characteristics are presented in Table 1.

**Table 1:**
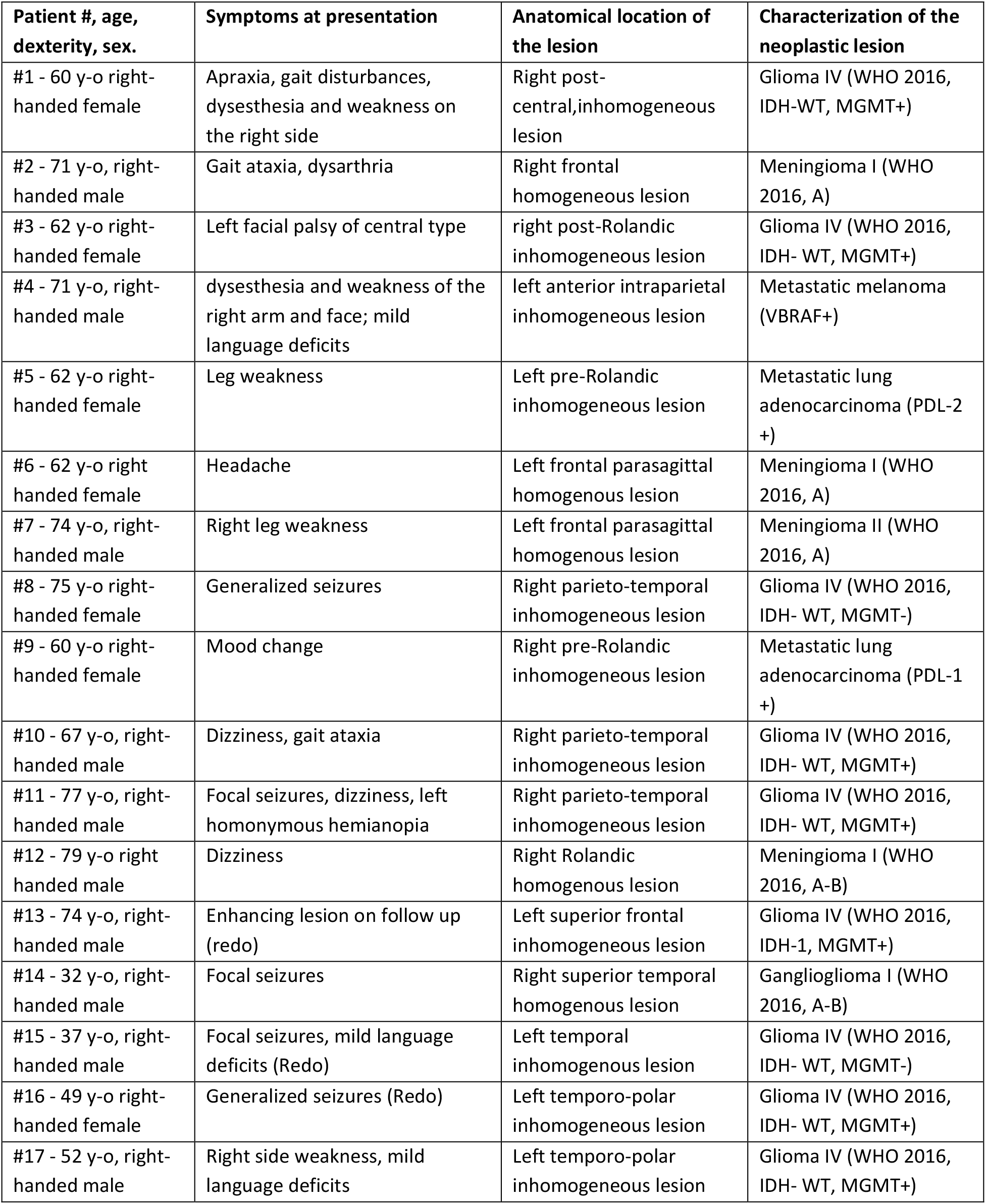
Demographic information on the group of patients.

### Stereotaxic neuronavigation and electrode placement

MRI scans of each patient’s brain were acquired before surgery on a 3T scanner with an eight-channel head coil (Signa *3T, General Electric* Healthcare, Milwaukee, USA). T1-weighted 3D MPRAGE images were acquired using the following parameters (echo train length: 1, TE: 2.67ms, TR: 2.000, matrix size: 256 × 246, slice thickness: 1mm). T2-weighted, FLAIR images were also acquired (TR 6000ms, TE 150mss, TI 2000ms). The reconstruction of the individual cortical surface was performed using Brainsuite (Brainsuite, UCLA Brain Mapping Center, San Francisco, USA “Shattuck DW and Leahy RM (2002)”). For a clearer intraoperative visualization of sulcal anatomy, a skull stripped T1 using a non-uniformity correction (Brainsuite, UCLA Brain Mapping Center, San Francisco, USA “Shattuck DW and Leahy RM (2002)”) or a FLAIR images was added to the 3D visualization of the Neuronavigation system (Stealth Station 7, Medtronic, Minneapolis, USA). Correspondence of 3D reconstruction and individual patient’s sulcal anatomy was then performed using the Neuronavigation pointer. Brain anatomy was systematically analysed prior to surgery so that main sulcal patterns of the postcentral and parietal region could be readily identified during actual surgery. The stimulating strip was placed parallel to the central sulcus, similarly to the montage used for studies of intra-operative motor evoked potentials, after identification of the central sulcus by phase reversal (Romstöck *et al*., 2002). Placement of the conditioning electrode strip was roughly planned a priori but was systematically reprogrammed when in presence of contingent surgical conditions preventing the placement of the strip in the desired position, such as presence of large vessels or space requirements by the ongoing surgical procedures.

### Anaesthesia and conventional IONM

The anaesthesia protocol applied was Total Intravenous Anaesthesia (TIVA). More precisely, a continuous infusion of Propofol (100-150 μg/kg/min) and Fentanyl (1μg/kg/min) was used, avoiding bolus. Short acting relaxants were administered for intubation purpose only and then avoided. Halogenated anaesthetic agents were never used. Since all patients were candidates for IONM of the corticospinal tract, standard neurophysiological monitoring and mapping was performed. This involved simultaneous acquisition of continuous electroencephalography (EEG), electrocorticography (ECoG), recording of free-running electromyographic (EMG) activity (ISIS-IOM, Inomed Medizintechnik GmbH, Emmendingen, Germany). Muscle MEPs were initially elicited by Transcranial Electrical Stimulation (TES) via corkscrew-like electrodes (Ambu^®^ Neuroline Corkscrew, Ambu, Copenhagen, Denmark) from the scalp. Short trains of 5 square-wave stimuli of 0.5 ms duration, and interstimulus interval (ISI) of 4ms were applied at a repetition rate up to 2 Hz through electrodes placed at C1 and C2 scalp sites, according to the 10/20 EEG system. Cortical and subcortical stimulation were performed using a monopolar probe (45 mm, angled 30°, Inomed Medizintechnik GmbH, Emmendingen, Germany) referenced to Fz. Stimulation parameters were as follows: a short train of five pulses, pulse duration 0.5 milliseconds; interstimulus interval (ISI) 2 ms at 1 Hz repetition rate. Cortical stimulation was anodal while subcortical stimulation was cathodal. Once the dura was opened, MEP monitoring was performed using a 6-contacts strip electrode (diameter 2.5 mm, space 10 mm, contact strips: 0.7 mm thin, 10 mm width, Inomed Medizintechnik GmbH, Emmendingen, Germany). EMG recordings were performed in a belly-tendon montage, by means of subcutaneous needle monopolar electrodes (Ambu^®^ Neuroline Subdermal, Ambu, Copenhagen, Denmark). The *orbicularis oris*, the ABP, the *biceps*, the *abductor hallucis* and the *tibialis anterior* muscles contralateral to the stimulated hemisphere were recorded.

### Dual strip stimulation

Direct electrical cortical stimulation was applied to the precentral gyrus (test stimuli) via a 6-contacts strip electrode and to the parietal cortex by means of a 6-contacts or an 8-contacts strip electrode (Figure 1 shows a schematic of the dual strip protocol and an example of surgical scenario). To optimize timing precision between the conditioning and the test stimuli, the conditioning stimuli were always delivered in a short train of 2 stimuli at 250 Hz and of 0.5 ms duration. Test stimuli were delivered with trains of the minimal duration required to elicit a stable MEP in the ABP. This results in test stimulation with one single stimulus in 1 patient, with 2 stimuli in 12 patients and with 3 stimuli in 3 patients. Intensity of test stimulation was set to obtain a MEP from the thenar muscle of around 500 uV peak-peak amplitude. The ISI was considered as the interval between the last stimulus of the conditioning train and the last pulse of the test train. The ISIs of 7, 13 and 18 ms were systematically explored in separate blocks for each of the test-stimulus electrodes. Every block contained at least 15 repetitions of the same dual stimulation. Dual-stimulation blocks were alternated with blocks with only test-stimuli so that the unconditioned MEP amplitude was monitored throughout the recording session. The timing of dual stimuli was manged entirely by the commercially available ISIS-IOM system (Inomed Medizintechnik GmbH, Emmendingen, Germany) by means of the “facilitation” function, that allows independent electrical stimulation through two separate output channels.

**Figure 1:**
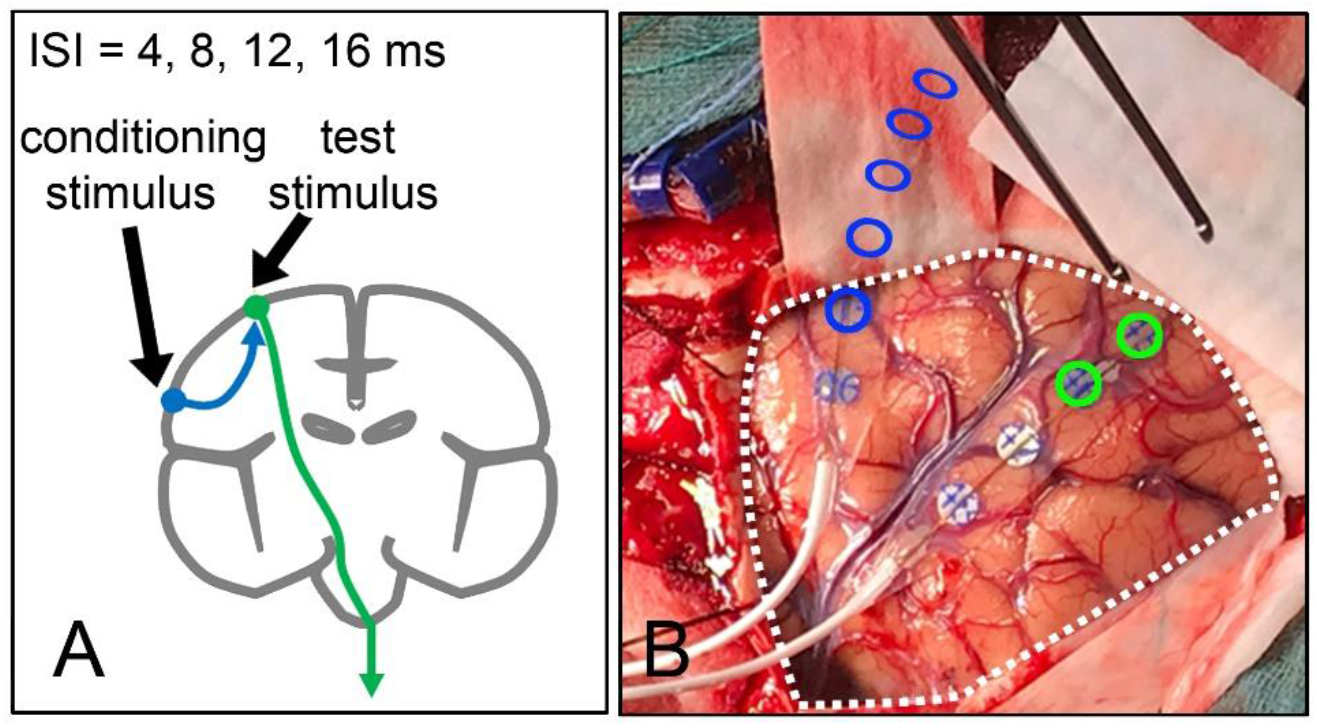
**A**-schematic representation of the dual strip protocol. Test stimuli are delivered to the motor cortex and produce a measurable motor evoked potential via the corticospinal tract (green arrow). In some trials the test stimulus is preceded by a conditioning stimulus, which alone cannot activate the corticospinal tract, applied to the parietal cortex (blue arrow). The inter-stimulus interval (ISI) ranges between 4 and 16 ms. The finding of MEPs to conditioning stimuli having a different amplitude or area than MEPs to test stimuli alone is considered evidence for functional connectivity between the two stimulated regions. The short duration of the ISI, in the range of milliseconds, indicates mono-or oligo-synaptic connections. **B**-Actual surgical scenario (patient #4). The dashed white line represents the borders of the craniotomy. The two stimulation strips are shown. The test stimulus dipole is indicated with the two green circles and is used to activate the corticospinal tract. The conditioning strip is placed over the parietal lobe. The dipoles used for conditioning stimuli are indicated with blue circles. Note that the electrode strip is inserted under the dura, therefore the electrode position extends well beyond the craniotomy.

### Data analysis

Pre-processing required the data to be exported in digital format and analysed with the MATLAB software. The EMG traces were band-pass filtered (50-2000 Hz) and rectified. The duration of the MEP was determined individually, and the corresponding area of the EMG recording was extracted. In this way each trial was characterized by a single number, i.e. the MEP area. We then proceeded to normalizing conditioned MEPs to test MEPs. However, MEPs to test stimuli alone are not stable throughout the surgical procedure because of strip movements. To correct for such variability, we normalized blocks of conditioned MEPs only to a sliding window of the blocks of test stimuli adjacent to each conditioned block. This was done by dividing the single conditioned MEP areas by the median of the test MEP areas. The resulting normalized conditioned MEP areas (normMEP) were used as main experimental variable. The main analysis was carried out in single patients, comparing normalized conditioned MEP areas from test stimuli by means of independent-samples t-tests. This test informs us whether, in single subjects, the trials in a given conditioned set are different from those in the test set and the direction of the change (excitation or inhibition). Significance threshold was corrected to account for the repeated comparisons. Analysis were performed therefore on single subjects, using a univariate approach. The results of single t-tests were therefore corrected for multiple comparisons in each participant. For example, patient #1 was tested on 20 cortical spots and therefore critical p-value was set to p=0.05/20 i.e. p=0.0025. Qualitative assessment of the effects of conditioning stimuli at the population level was performed by plotting on a standardized surface map of the parietal cortex the sites of conditioning stimulation of all participants, indicating electrodes that had a significant effect on test stimuli in single participants. To this purpose, we used the frameless stereotaxic neuronavigation system to pinpoint ion individual brain anatomies the real position of strip electrodes and the trajectory of strips that were not readily visible because in the subdural space. The parietal region shows a considerable inter-individual variability, therefore, to remap individual anatomies on the standardized space we made reference to the anatomical study of Zlatkina and Petrides (2014), which resulted in slight warping of the strip positions. Figure 2 indicates individual brain anatomies of the 16 patients together with the conditioning strip electrodes and Figure 3 Indicates the population data in the standardized space.

**Figure 2:**
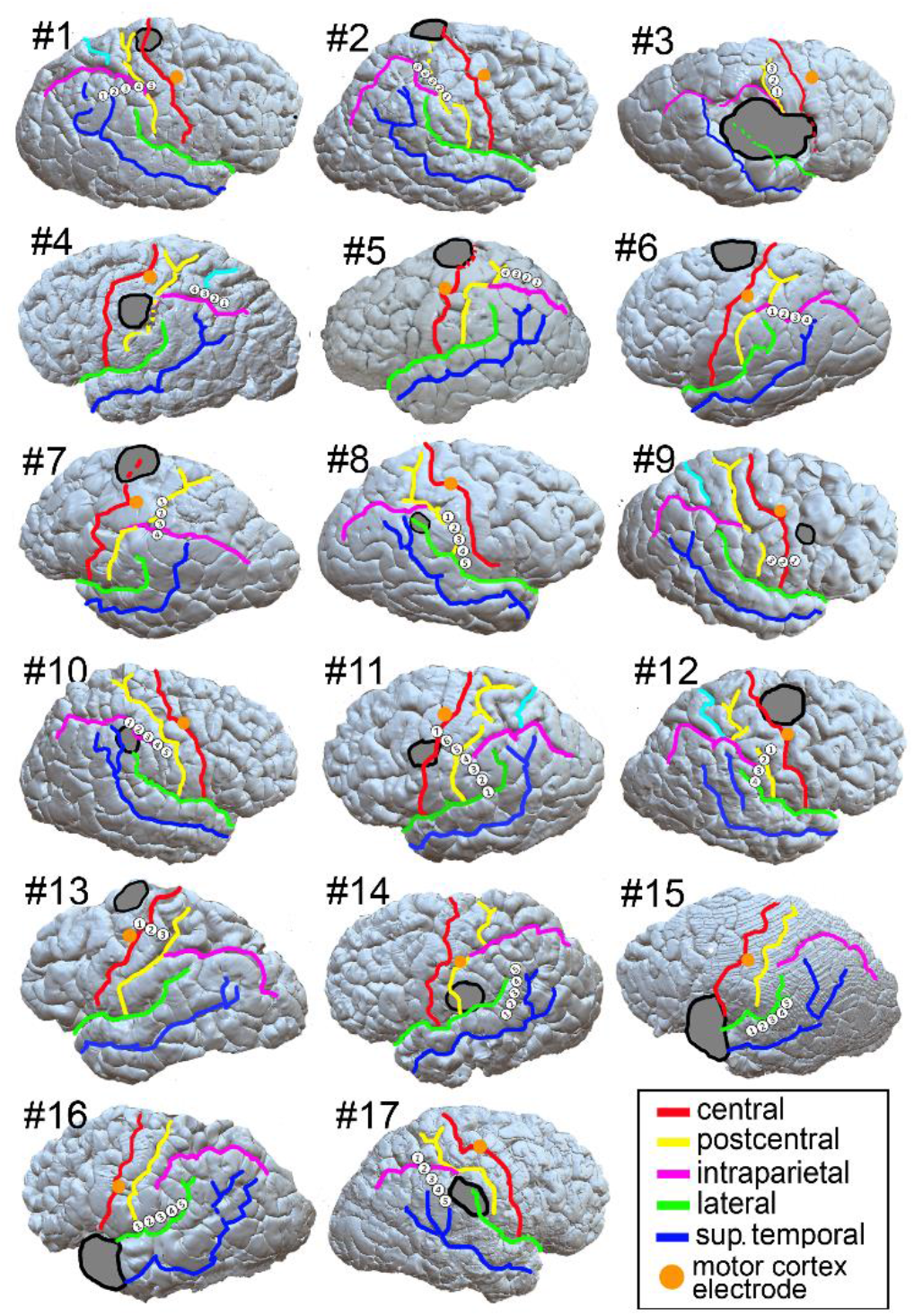
rendering of the individual brains (only the stimulated hemisphere is shown. The projection of the lesion on the surface is indicated with the grey shape. Main cortical sulci are indicated with a colour code as indicated in the legend. The orange spot indicates the point of corticospinal stimulation (test stimulus). The white numbered circles indicate the position of the cathode of conditioning stimulation. (note that conditioning stimuli have been delivered in bipolar modality).

**Figure 3:**
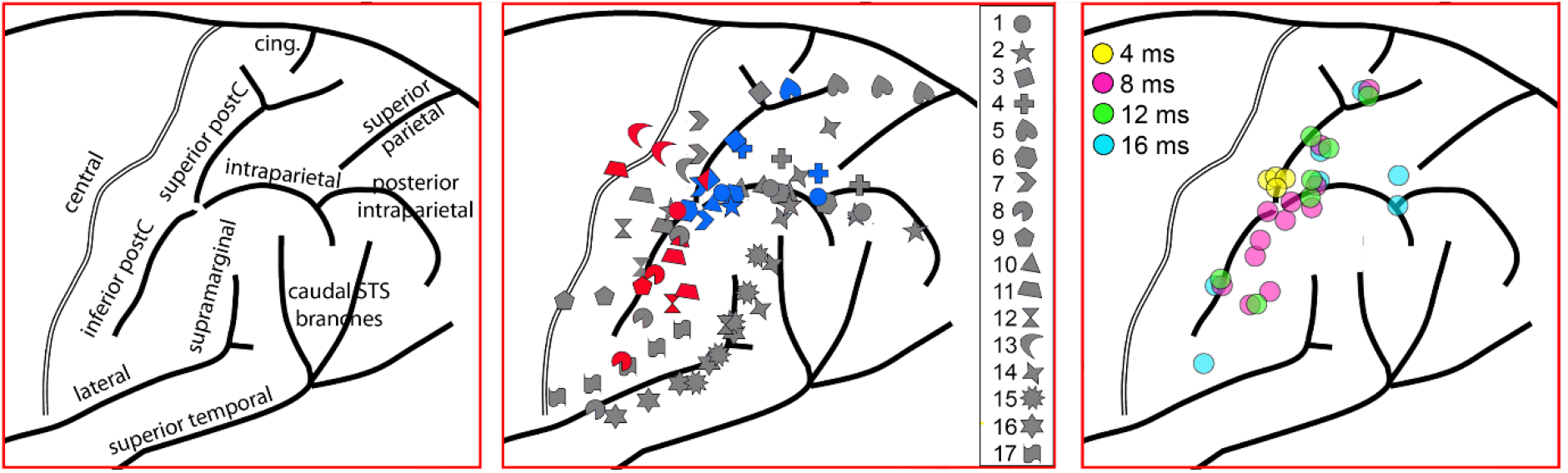
standardized parietal anatomy showing the location of each participant’s conditioning electrodes in the parietal cortex. The anatomical template is based on (Zlatkina and Petrides, 2014; Zlatkina et al., 2016). The **left panel** shows the label of the main sulci. The **middle panel** indicates the positions of conditioning stimuli (cathodes) in each patient. The legend links symbols to each patient’s numbers. Grey-filled symbols indicate spots with no significant effect. Blue-filled symbols indicate spots with significant inhibitory conditioning effects. Red-filled symbols indicate spots with significant excitatory conditioning effects. Spots in which both excitatory and inhibitory effects were observed at different ISIs are indicated with both red and blue filling. The **right panel** shows only conditioning spots with significant effects (active spots), pooled across patients and grouped according to the ISI at which an effect was observed.

## Results

In all participants it was possible to stimulate at least one conditioning spot, with a variable number of 3-6. Figure 2 shows each patient’s anatomy together with lesion location and electrode placement. For the sake of clarity, participants have been numbered according to the presence of excitatory effect, inhibitory effects or no effect. We observed in most subjects a significant modulation of MEPs by conditioning stimuli in one specific electrode, at specific timings. The individual results are reported in Figures 3 and 4 and in Table 2. We observed both inhibitory and excitatory effects of conditioning stimuli at different ISIs. The systematic between-subject variations inherent in the mapping technique were reflected in the variability of the ISIs at which conditioning stimuli exerted a significant effect on corticospinal excitability which ranged from 4ms to 16 ms. 6 participants showed only inhibitory effects, 3 showed mixed effects, 4 showed faciliatory effects and 4 did not show any effect of conditioning stimuli on corticospinal excitability.

**Figure 4:**
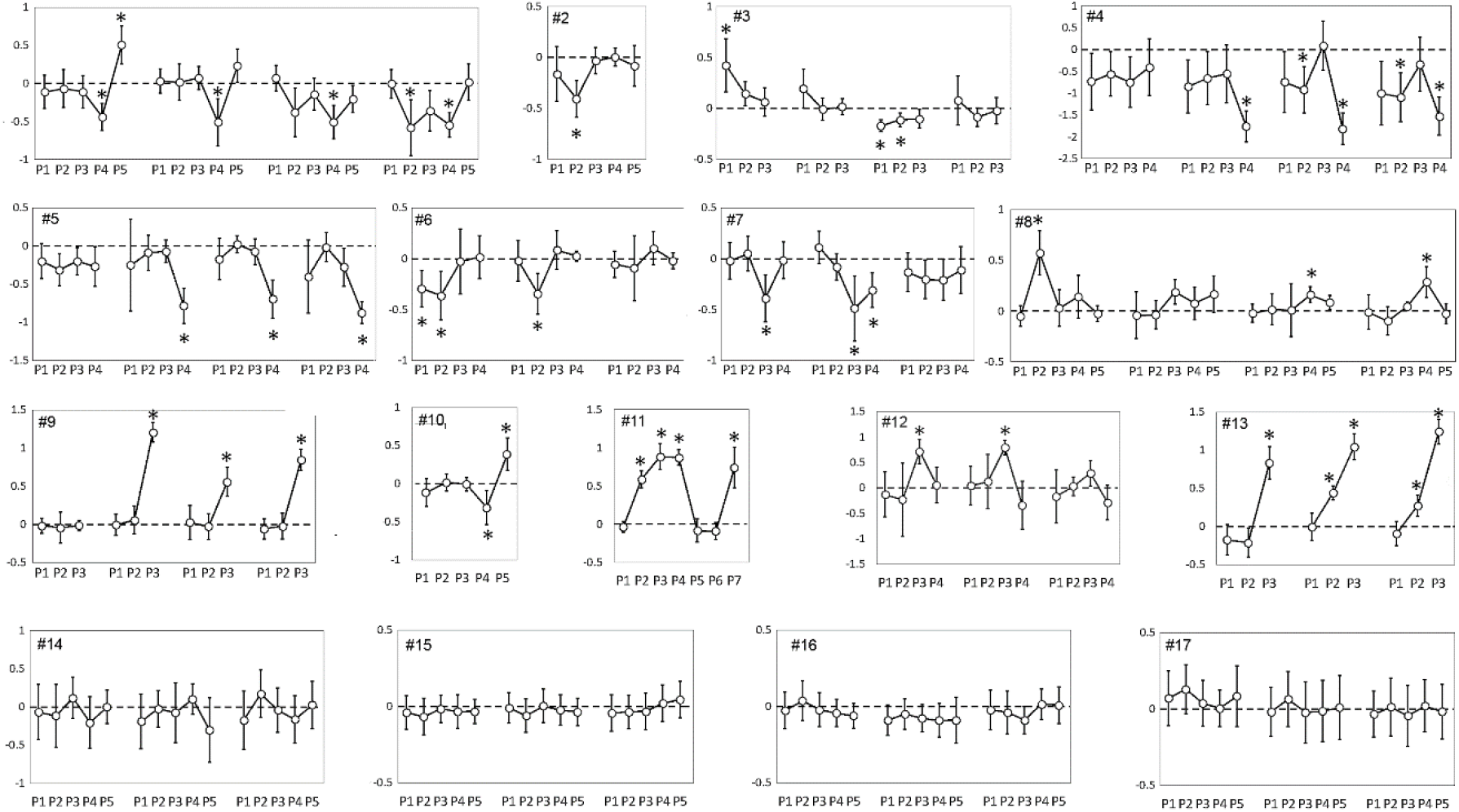
Comprehensive data from all patients. The plots indicate the mean value of the log-transformed ratios between conditioned and unconditioned MEPs. Negative values indicate that the mean conditioned MEP is smaller than the mean test MEP (inhibitory effects). Positive values indicate that mean conditioned MEPs are larger than mean test MEPs (excitatory effects). All log-ratios have been tested by single-sample t-tests against a null hypothesis of mean value = 0. Significance threshold was Bonferroni corrected. Asterisks indicate significant comparisons. Error bars indicate 95% confidence intervals of the mean).

**Table 2:** statistics on individual patients, for each conditioning electrode and each ISI. The data reported correspond to the log of the ratio between conditioned MEPs and test MEPs. The table reports the mean value (standard deviation), and the t-statistics of the single-sample t-tests: t-value (degrees of freedom) and p-value. Please note that the log-ratio has no dimensions because it is obtained by dividing two identical dimensions. Note also that degrees of freedom are variable according to the number of single trials that were performed in that particular condition. Conditions with p-values exceeding the Bonferroni-corrected significance threshold are highlighted in bold. A corresponding graphical representation of the data is provided in Figure 4.

| patient | ISI: | conditioning stimulus cathode |  |  |  |  |
| --- | --- | --- | --- | --- | --- | --- |
|  |  | P1 | P2 | P3 | P4 | P5 |
| #1 | 4ms | -0.113 (sd: 0.22)<br>t(19)=-1.17<br>p=0.2582 | -0.069 (sd: 0.25)<br>t(19)=-0.63<br>p=0.5378 | -0.112 (sd: 0.21)<br>t(19)=-1.18<br>p=0.2544 | **-0.441 (sd: 0.18)**<br>**t(19)=-5.62**<br>**p<0.00001** | **0.508 (sd: 0.25)**<br>**t(19)=4.59**<br>**p=0.0002** |
|  | 8ms | 0.03 (sd: 0.16)<br>t(19)=0.42<br>p=0.6856 | 0.016 (sd: 0.24)<br>t(19)=0.15<br>p=0.8814 | 0.071 (sd: 0.15)<br>t(19)=1.08<br>p=0.2949 | **-0.513 (sd: 0.31)**<br>**t(19)=-3.69**<br>**p=0.0016** | 0.232 (sd: 0.22)<br>t(19)=2.36<br>p=0.0288 |
|  | 12ms | 0.068 (sd: 0.17)<br>t(19)=0.92<br>p=0.3688 | -0.38 (sd: 0.32)<br>t(19)=-2.66<br>p=0.0156 | -0.143 (sd: 0.21)<br>t(19)=-1.54<br>p=0.1407 | **-0.511 (sd: 0.22)**<br>**t(19)=-5.16**<br>**p=0.0001** | -0.209 (sd: 0.18)<br>t(19)=-2.53<br>p=0.0201 |
|  | 16ms | -0.005 (sd: 0.19)<br>t(19)=-0.06<br>p=0.9584 | **-0.585 (sd: 0.37)**<br>**t(19)=-3.51**<br>**p=0.0023** | -0.362 (sd: 0.27)<br>t(19)=-2.98<br>p=0.0077 | **-0.549 (sd: 0.16)**<br>**t(19)=-7.81**<br>**p<0.00001** | 0.016 (sd: 0.24)<br>t(19)=0.15<br>p=0.8814 |
| #2 | 8ms | -0.164 (sd: 0.27)<br>t(14)=-1.39<br>p=0.19 | **-0.408 (sd: 0.18)**<br>**t(14)=-5.02**<br>**p=0.0002** | -0.035 (sd: 0.13)<br>t(14)=-0.6<br>p=0.57 | 0.002 (sd: 0.09)<br>t(14)=0.05<br>p=0.96 | -0.084 (sd: 0.2)<br>t(14)=-0.93<br>p=0.37 |
| #3 | 4ms | **0.421 (sd: 0.26)**<br>**t(14)=3.59**<br>**p=0.003** | 0.142 (sd: 0.12)<br>t(14)=2.68<br>p=0.018 | 0.062 (sd: 0.14)<br>t(14)=0.96<br>p=0.35 |  |  |
|  | 8ms | 0.193 (sd: 0.19)<br>t(14)=2.32<br>p=0.03 | -0.01 (sd: 0.11)<br>t(14)=-0.2<br>p=0.84 | 0.015 (sd: 0.08)<br>t(14)=0.44<br>p=0.66 |  |  |
|  | 12ms | **-0.172 (sd: 0.06)**<br>**t(14)=-6.26**<br>**p<0.00001** | **-0.115 (sd: 0.07)**<br>**t(14)=-3.77**<br>**p=0.002** | -0.104 (sd: 0.09)<br>t(14)=-2.57<br>p=0.02 |  |  |
|  | 16ms | 0.077 (sd: 0.24)<br>t(14)=0.72<br>p=0.48 | -0.087 (sd: 0.09)<br>t(14)=-2.13<br>p=0.05 | -0.023 (sd: 0.13)<br>t(14)=-0.41<br>p=0.69 |  |  |
| #4 | 4ms | -0.732 (sd: 0.56)<br>t(14)=-2.53<br>p=0.02 | -0.56 (sd: 0.44)<br>t(14)=-2.45<br>p=0.028 | -0.747 (sd: 0.5)<br>t(14)=-2.88<br>p=0.01 | -0.407 (sd: 0.56)<br>t(14)=-1.41<br>p=0.18 |  |
|  | 8ms | -0.85 (sd: 0.53)<br>t(14)=-3.12<br>p=0.008 | -0.653 (sd: 0.53)<br>t(14)=-2.38<br>p=0.032 | -0.556 (sd: 0.57)<br>t(14)=-1.9<br>p=0.078 | **-1.762 (sd: 0.3)**<br>**t(14)=-11.33**<br>**p<0.00001** |  |
|  | 12ms | -0.744 (sd: 0.6)<br>t(14)=-2.4<br>p=0.03 | -0.92 (sd: 0.46)<br>t(14)=-3.88<br>p=0.002 | 0.09 (sd: 0.49)<br>t(14)=0.36<br>p=0.73 | **-1.821 (sd: 0.31)**<br>**t(14)=-11.27**<br>**p<0.00001** |  |
|  | 16ms | -1.001 (sd: 0.63)<br>t(14)=-3.09<br>p=0.008 | **-1.093 (sd: 0.49)**<br>**t(14)=-4.36**<br>**p=0.0007** | -0.334 (sd: 0.54)<br>t(14)=-1.21<br>p=0.25 | **-1.529 (sd: 0.38)**<br>**t(14)=-7.81**<br>**p<0.00001** |  |
| #5 | 4ms | -0.201 (sd: 0.23)<br>t(19)=-1.92<br>p=0.069 | -0.313 (sd: 0.21)<br>t(19)=-3.29<br>p=0.0038 | -0.2 (sd: 0.18)<br>t(19)=-2.43<br>p=0.02 | -0.269 (sd: 0.26)<br>t(19)=-2.34<br>p=0.03 |  |
|  | 8ms | -0.251 (sd: 0.6)<br>t(19)=-0.93<br>p=0.36 | -0.087 (sd: 0.23)<br>t(19)=-0.83<br>p=0.41 | -0.07 (sd: 0.15)<br>t(19)=-1.02<br>p=0.32 | **-0.783 (sd: 0.23)**<br>**t(19)=-7.75**<br>**p<0.00001** |  |
|  | 12ms | -0.173 (sd: 0.27)<br>t(19)=-1.43<br>p=0.16 | 0.023 (sd: 0.11)<br>t(19)=0.47<br>p=0.65 | -0.076 (sd: 0.17)<br>t(19)=-1.01<br>p=0.33 | **-0.696 (sd: 0.25)**<br>**t(19)=-6.23**<br>**p<0.00001** |  |
|  | 16ms | -0.403 (sd: 0.48)<br>t(19)=-1.88<br>p=0.07 | -0.018 (sd: 0.19)<br>t(19)=-0.21<br>p=0.83 | -0.276 (sd: 0.25)<br>t(19)=-2.51<br>p=0.022 | **-0.874 (sd: 0.14)**<br>**t(19)=-13.62**<br>**p<0.00001** |  |
| #6 | 8ms | **-0.291 (sd: 0.18)**<br>**t(12)=-3.59**<br>**p=0.004** | -0.362 (sd: 0.24)<br>t(12)=-3.36<br>p=0.006 | -0.027 (sd: 0.32)<br>t(12)=-0.19<br>p=0.86 | 0.016 (sd: 0.21)<br>t(12)=0.17<br>p=0.87 |  |
|  | 12ms | -0.018 (sd: 0.2)<br>t(12)=-0.2<br>p=0.84 | **-0.343 (sd: 0.2)**<br>**t(12)=-3.79**<br>**p=0.0026** | 0.086 (sd: 0.19)<br>t(12)=1.02<br>p=0.33 | 0.026 (sd: 0.05)<br>t(12)=1.13<br>p=0.29 |  |
|  | 16ms | -0.054 (sd: 0.13)<br>t(12)=-0.93<br>p=0.37 | -0.093 (sd: 0.32)<br>t(12)=-0.65<br>p=0.53 | 0.104 (sd: 0.16)<br>t(12)=1.48<br>p=0.17 | -0.019 (sd: 0.08)<br>t(12)=-0.52<br>p=0.61 |  |
| #7 | 4ms | -0.02 (sd: 0.18)<br>t(15)=-0.25<br>p=0.80 | 0.049 (sd: 0.17)<br>t(15)=0.65<br>p=0.52 | **-0.388 (sd: 0.23)**<br>**t(15)=-3.71**<br>**p=0.002** | -0.015 (sd: 0.18)<br>t(15)=-0.19<br>p=0.85 |  |
|  | 8ms | 0.113 (sd: 0.16)<br>t(15)=1.55<br>p=0.14 | -0.081 (sd: 0.13)<br>t(15)=-1.36<br>p=0.20 | -0.488 (sd: 0.32)<br>t(15)=-3.42<br>p=0.004 | **-0.309 (sd: 0.17)**<br>**t(15)=-4.1**<br>**p=0.0009** |  |
|  | 16ms | -0.132 (sd: 0.19)<br>t(15)=-1.55<br>p=0.14 | -0.203 (sd: 0.19)<br>t(15)=-2.44<br>p=0.03 | -0.209 (sd: 0.2)<br>t(15)=-2.39<br>p=0.030 | -0.111 (sd: 0.23)<br>t(15)=-1.1<br>p=0.29 |  |
| #8 | 4ms | -0.051 (sd: 0.1)<br>t(14)=-1.16<br>p=0.27 | **0.573 (sd: 0.22)**<br>**t(14)=5.72**<br>**p=0.0001** | 0.027 (sd: 0.18)<br>t(14)=0.34<br>p=0.74 | 0.141 (sd: 0.21)<br>t(14)=1.47<br>p=0.16 | -0.027 (sd: 0.08)<br>t(14)=-0.74<br>p=0.48 |
|  | 8ms | -0.043 (sd: 0.23)<br>t(14)=-0.43<br>p=0.67 | -0.039 (sd: 0.14)<br>t(14)=-0.65<br>p=0.53 | 0.188 (sd: 0.12)<br>t(14)=3.37<br>p=0.004 | 0.076 (sd: 0.16)<br>t(14)=1.05<br>p=0.31 | 0.165 (sd: 0.18)<br>t(14)=2.05<br>p=0.06 |
|  | 12ms | -0.022 (sd: 0.09)<br>t(14)=-0.54<br>p=0.60 | 0.016 (sd: 0.15)<br>t(14)=0.23<br>p=0.82 | 0.007 (sd: 0.26)<br>t(14)=0.06<br>p=0.95 | **0.16 (sd: 0.08)**<br>**t(14)=4.49**<br>**p=0.0005** | 0.084 (sd: 0.07)<br>t(14)=2.54<br>p=0.02 |
|  | 16ms | -0.012 (sd: 0.17)<br>t(14)=-0.15<br>p=0.88 | -0.099 (sd: 0.14)<br>t(14)=-1.6<br>p=0.13 | 0.046 (sd: 0.04)<br>t(14)=2.93<br>p=0.012 | **0.285 (sd: 0.15)**<br>**t(14)=4.21**<br>**p=0.0009** | -0.026 (sd: 0.1)<br>t(14)=-0.57<br>p=0.58 |
| #9 | 4ms | -0.018 (sd: 0.1)<br>t(22)=-0.39<br>p=0.70 | -0.042 (sd: 0.2)<br>t(22)=-0.46<br>p=0.65 | -0.014 (sd: 0.07)<br>t(22)=-0.47<br>p=0.65 |  |  |
|  | 8ms | -0.006 (sd: 0.14)<br>t(22)=-0.09<br>p=0.93 | 0.059 (sd: 0.18)<br>t(22)=0.72<br>p=0.48 | **1.208 (sd: 0.13)**<br>**t(22)=20.46**<br>**p<0.00001** |  |  |
|  | 12ms | 0.026 (sd: 0.22)<br>t(22)=0.26<br>p=0.7932 | -0.026 (sd: 0.17)<br>t(22)=-0.34<br>p=0.73 | **0.558 (sd: 0.19)**<br>**t(22)=6.62**<br>**p<0.00001** |  |  |
|  | 16ms | -0.057 (sd: 0.13)<br>t(22)=-0.95<br>p=0.35 | -0.023 (sd: 0.17)<br>t(22)=-0.3<br>p=0.77 | **0.845 (sd: 0.14)**<br>**t(22)=13.17**<br>**p<0.00001** |  |  |
| #10 | 8ms | -0.116 (sd: 0.18)<br>t(13)=-1.46<br>p=0.1676 | 0.016 (sd: 0.11)<br>t(13)=0.33<br>p=0.7491 | -0.007 (sd: 0.09)<br>t(13)=-0.17<br>p=0.8681 | **-0.312 (sd: 0.22)**<br>**t(13)=-3.17**<br>**p=0.0075** | **0.387 (sd: 0.21)**<br>**t(13)=4.12**<br>**p=0.0012** |
| #11 | 8ms | **0.885 (sd: 0.17)**<br>**t(13)=11.58**<br>**p<0.00001** | **0.871 (sd: 0.1)**<br>**t(13)=19.65**<br>**p<0.00001** | -0.085 (sd: 0.15)<br>t(13)=-1.3<br>p=0.22 | -0.091 (sd: 0.11)<br>t(13)=-1.87<br>p=0.084 | **0.74 (sd: 0.27)**<br>**t(13)=6.09**<br>**p<0.00001** |
| #12 | 8ms | -0.131 (sd: 0.44)<br>t(15)=-0.67<br>p=0.51 | -0.231 (sd: 0.72)<br>t(15)=-0.72<br>p=0.49 | **0.715 (sd: 0.24)**<br>**t(15)=6.81**<br>**p<0.00001** | 0.058 (sd: 0.35)<br>t(15)=0.37<br>p=0.71 |  |
|  | 12ms | 0.041 (sd: 0.38)<br>t(15)=0.24<br>p=0.81 | 0.122 (sd: 0.53)<br>t(15)=0.52<br>p=0.61 | **0.791 (sd: 0.14)**<br>**t(15)=12.91**<br>**p<0.00001** | -0.343 (sd: 0.47)<br>t(15)=-1.64<br>p=0.12 |  |
|  | 16ms | -0.167 (sd: 0.52)<br>t(15)=-0.72<br>p=0.48 | 0.03 (sd: 0.18)<br>t(15)=0.37<br>p=0.71 | 0.288 (sd: 0.25)<br>t(15)=2.56<br>p=0.02 | -0.288 (sd: 0.34)<br>t(15)=-1.9<br>p=0.08 |  |
| #13 | 8ms | -0.174 (sd: 0.2)<br>t(14)=-1.92<br>p=0.07 | -0.211 (sd: 0.19)<br>t(14)=-2.46<br>p=0.03 | **0.833 (sd: 0.21)**<br>**t(14)=8.93**<br>**p<0.00001** |  |  |
|  | 12ms | -0.005 (sd: 0.18)<br>t(14)=-0.06<br>p=0.95 | **0.441 (sd: 0.09)**<br>**t(14)=10.64**<br>**p<0.00001** | **1.044 (sd: 0.17)**<br>**t(14)=14.06**<br>**p<0.00001** |  |  |
|  | 16ms | -0.093 (sd: 0.16)<br>t(14)=-1.29<br>p=0.22 | **0.274 (sd: 0.14)**<br>**t(14)=4.54**<br>**p=0.0005** | **1.242 (sd: 0.16)**<br>**t(14)=17.91**<br>**p<0.00001** |  |  |
| #14 | 8ms | -0.039 (sd: 0.11)<br>t(13)=-0.81<br>p=0.43 | -0.068 (sd: 0.12)<br>t(13)=-1.29<br>p=0.22 | -0.015 (sd: 0.09)<br>t(13)=-0.37<br>p=0.72 | -0.034 (sd: 0.11)<br>t(13)=-0.7<br>p=0.49 | -0.032 (sd: 0.08)<br>t(13)=-0.91<br>p=0.38 |
|  | 12ms | -0.009 (sd: 0.1)<br>t(13)=-0.2<br>p=0.85 | -0.061 (sd: 0.11)<br>t(13)=-1.24<br>p=0.24 | 0.004 (sd: 0.11)<br>t(13)=0.08<br>p=0.94 | -0.023 (sd: 0.1)<br>t(13)=-0.53<br>p=0.61 | -0.035 (sd: 0.09)<br>t(13)=-0.87<br>p=0.40 |
|  | 16ms | -0.043 (sd: 0.12)<br>t(13)=-0.81<br>p=0.43 | -0.036 (sd: 0.11)<br>t(13)=-0.73<br>p=0.48 | -0.032 (sd: 0.12)<br>t(13)=-0.6<br>p=0.56 | 0.02 (sd: 0.12)<br>t(13)=0.39<br>p=0.71 | 0.046 (sd: 0.12)<br>t(13)=0.87<br>p=0.40 |
| #15 | 8ms | -0.026 (sd: 0.12)<br>t(13)=-0.48<br>p=0.64 | 0.037 (sd: 0.13)<br>t(13)=0.63<br>p=0.54 | -0.021 (sd: 0.11)<br>t(13)=-0.43<br>p=0.67 | -0.043 (sd: 0.09)<br>t(13)=-1.12<br>p=0.28 | -0.061 (sd: 0.08)<br>t(13)=-1.74<br>p=0.11 |
|  | 12ms | -0.091 (sd: 0.1)<br>t(13)=-2.07<br>p=0.06 | -0.049 (sd: 0.1)<br>t(13)=-1.06<br>p=0.31 | -0.077 (sd: 0.09)<br>t(13)=-1.86<br>p=0.09 | -0.091 (sd: 0.11)<br>t(13)=-1.85<br>p=0.08 | -0.089 (sd: 0.15)<br>t(13)=-1.37<br>p=0.20 |
|  | 16ms | -0.021 (sd: 0.13)<br>t(13)=-0.37<br>p=0.72 | -0.038 (sd: 0.14)<br>t(13)=-0.59<br>p=0.56 | -0.09 (sd: 0.09)<br>t(13)=-2.13<br>p=0.05 | 0.015 (sd: 0.1)<br>t(13)=0.35<br>p=0.72 | 0.008 (sd: 0.12)<br>t(13)=0.15<br>p=0.89 |
| #16 | 8ms | 0.071 (sd: 0.18)<br>t(17)=0.9<br>p=0.38 | 0.13 (sd: 0.16)<br>t(17)=1.88<br>p=0.07 | 0.038 (sd: 0.15)<br>t(17)=0.57<br>p=0.58 | 0.005 (sd: 0.12)<br>t(17)=0.1<br>p=0.93 | 0.084 (sd: 0.2)<br>t(17)=0.94<br>p=0.36 |
|  | 12ms | -0.018 (sd: 0.16)<br>t(17)=-0.26<br>p=0.80 | 0.065 (sd: 0.18)<br>t(17)=0.82<br>p=0.42 | -0.022 (sd: 0.2)<br>t(17)=-0.25<br>p=0.80 | -0.013 (sd: 0.2)<br>t(17)=-0.15<br>p=0.88 | 0.01 (sd: 0.21)<br>t(17)=0.11<br>p=0.91 |
|  | 16ms | -0.034 (sd: 0.15)<br>t(-1)=-0.51<br>p=0.61 | 0.013 (sd: 0.19)<br>t(17)=0.15<br>p=0.88 | -0.044 (sd: 0.2)<br>t(17)=-0.49<br>p=0.63 | 0.021 (sd: 0.17)<br>t(17)=0.28<br>p=0.78 | -0.017 (sd: 0.18)<br>t(17)=-0.22<br>p=0.83 |
| #17 | 4ms | -0.066 (sd: 0.36)<br>t(13)=-0.42<br>p=0.68 | -0.116 (sd: 0.41)<br>t(13)=-0.64<br>p=0.53 | 0.118 (sd: 0.27)<br>t(13)=0.97<br>p=0.35 | -0.205 (sd: 0.34)<br>t(13)=-1.37<br>p=0.19 | 0.002 (sd: 0.22)<br>t(13)=0.02<br>p=0.98 |
|  | 8ms | -0.189 (sd: 0.36)<br>t(13)=-1.18<br>p=0.26 | -0.022 (sd: 0.24)<br>t(13)=-0.2<br>p=0.85 | -0.074 (sd: 0.39)<br>t(13)=-0.43<br>p=0.67 | 0.106 (sd: 0.2)<br>t(13)=1.18<br>p=0.26 | -0.302 (sd: 0.42)<br>t(13)=-1.61<br>p=0.13 |
|  | 16ms | -0.173 (sd: 0.38)<br>t(13)=-1.03<br>p=0.32 | 0.174 (sd: 0.31)<br>t(13)=1.28<br>p=0.22 | -0.04 (sd: 0.29)<br>t(13)=-0.3<br>p=0.77 | -0.163 (sd: 0.31)<br>t(13)=-1.18<br>p=0.26 | 0.026 (sd: 0.31)<br>t(13)=0.19<br>p=0.85 |

### Anatomical localization and chronometry of conditioning stimulus effects

Each patient was stimulated with conditioning stimuli in 2-5 pairs of stimulating electrodes (Figure 2 indicates the cathode of the bipolar stimulation montages). Active spots were localized all along a region of the PPC immediately posterior to the post-central sulcus (Figure 3 – middle panel). In addition, in the few participants in which the conditioning stimulus strip reached the central sulcus, we observed a small cluster of active spots corresponding to the hand motor cortex. We did not observe significant effects from stimulation of the postcentral gyrus at the ISIs adopted here. The polarity of the effect was spatially organized. We observed inhibitory effects of conditioning stimuli applied to the superior parietal lobule and the anterior intraparietal region. We found excitatory effects from conditioning stimuli applied to the inferior parietal lobule. Most patients showed only facilitatory or inhibitory effects in all stimulation dipoles. In two patients (#1 and #10) we observed a change in polarity of the effect from inhibitory to facilitatory moving the stimulating electrode ventrally and rostrally. All spots showed the same polarity of effect, whenever present, at all ISIs, with the sole exception of a single spot in a single patient (#3), localized in the intraparietal region, that showed facilitatory effects at the 4ms ISI and inhibitory effects at the 12 ms ISI. Figure 3 – right panel, shows the anatomical localization of effective conditioning stimuli, grouped by ISI. The spots with effects at 4 ms were all clustered in proximity of the hand motor area, while effective spots at higher ISIs increased gradually their distance from hand-M1.

## Discussion

### Three distinct regions in the parietal cortex exert specific short-latency effects on upper-limb corticospinal excitability

In the present work we demonstrate for the first time the existence of direct parietal-motor functional connections in humans by means of direct cortical stimulation. The presence of short-latency modulations of conditioning stimuli implies that the two regions are functionally connected. (Koch and Rothwell, 2009). We identified several cortical spots in the posterior parietal cortex that exert a short-latency effect on the excitability of the corticospinal pathway to the upper limb. Combining spatial distribution and polarity (excitatory or inhibitory) of the conditioning effects, we identified three distinct regions: a ventral region, corresponding to the part of the supramarginal gyrus immediately posterior to the inferior postcentral sulcus, extending ventrally to the parietal opercular region, a dorsal inhibitory region comprising the junction between the intraparietal sulcus and the precentral sulcus and the portion of the superior parietal lobule adjacent to the postcentral sulcus. A third region was indicated by a small cluster of 2 electrodes along the intraparietal sulcus, around its middle portion. However, we should consider that the spatial sampling procedure employed here suffers from a main limitation, that is, conditioning stimuli have been delivered on the crown of the sulci, because the surgical procedures do not imply the opening of the arachnoid and widening of the sulci. As such, our map of the parietal cortex is patchy and strongly biased towards the crown of the gyri.

### Widespread representation of upper limb movements in the posterior parietal cortex

In our study we focussed on the motor representation of the distal upper limb. A striking result is that the posterior parietal cortex seems to contain a widespread representation of the upper limb. Having tested only one effector, we cannot draw any conclusions on somatotopy, but our partial results argue against a possible somatotopic arrangement of motor representations in the PPC, in opposition of what we could have expected in the premotor region where rough somatotopy is suggested by several studies (Cunningham *et al*., 2013). Conversely, the posterior parietal cortex, albeit embedded with consistent motor representations, has not been shown to be organized effector-wise in humans. Action representations in the PPC of humans seem to show an upper limb preference and a spatial organization that reflects the type of action rather than the effector used, except for eye movements that are supported by a specialized network (Grefkes and Fink, 2005). Most authors seem to agree on a motor map of the rostral PPC organized in a medial-lateral system. Spatially-oriented stimuli are coded in the SPL, object-directed movements in the mid-portion, corresponding to the intraparietal sulcus and more complex hand actions such as symbolic movements and tool use are coded in the IPL (Gallivan and Culham, 2015; Orban, 2016). Consequently, the finding of hand representations throughout its medio-lateral extension is supported by current knowledge on the physiology of motor properties of the human PPC.

### A direct parieto-motor pathway in humans

Neuroimaging studies in humans however, are not able to specify the actual neural pathways by which the PPC can modulate movement. The novelty of the present study is providing compelling evidence that one possible neural substrate of the PPC influence on action is fast, probably direct, parieto-motor connectivity. The temporal characteristics of the parieto-motor interactions are illustrated in Figure 3C. We show a general pattern of active spots modulating the output of hand-M1 at ISIs that are roughly proportional to the distance between the active spot and M1, compatibly with axonal conduction of action potentials. Active spots that exert modulation at 4 ms ISIs are all clustered nearby the hand-M1. Active spots that are effective at longer ISIs are located progressively further away from the hand-M1. The two spots in the mid-intraparietal cluster both appeared at 16 ms. This pattern indicates that effective ISIs scale positively with linear distance to hand-M1. There are two main inferences to be made from this finding: First, that the effect on conditioning stimuli on the PPC is not likely to be due to current spread to M1, because current spread is quasi-instantaneous, and the latency of its effect would not increase with distance. Second, the increasing latency of the conditioning effects with increasing distance from M1 argues against the possibility that the site of interaction between parietal output and the corticospinal pathway is subcortical or spinal because in that case we would expect similar latencies of conditioning effects. On the contrary, the pattern of co-variation of distance to hand-M1 with effective ISI is strongly in favour of cortico-cortical connections between PPC and hand-M1 mediating the conditioning effect.

### Effects of anaesthesia

All our patients have been tested under the TIVA protocol. The main effect of propofol is a strong enhancement of GABAergic inputs (Franks, 2008). In terms of brain connectivity, Propofol dampens extensive cortico-cortical connections, according to TMS-EEG studies (Sarasso et al., 2015). The strength of oligosynaptic pathways is affected but generally not to the level of a conduction block, as witnessed by the validity of somatosensory evoked potentials and motor evoked potentials, which are mediated by multi-synaptic neural chains. The TIVA protocol is titrable, and it was systematically kept at low levels of neural suppression (see methods). Therefore, the anaesthesiologic setting employed here is appropriate to test mono- or oligo-synaptic connections, though we cannot make any inference on how these connections would work in the awake state. Summing up, the implications of testing cortico-cortical connectivity under anaesthesia are that: A) significant effects can be considered as genuine expression of oligo-synaptic connections, B) non-significant stimulations could underlie either no connections or multi-synaptic connections which are dampened by anaesthesia and C) we cannot make any inference on how the highlighted connections would function in the awake state and even more so during active tasks.

### Relation to connectivity data obtained with non-invasive brain stimulation

Ipsilateral PPC-M1 connections have been tested non-invasively by dual-coil TMS in awake subjects in a series of studies (Koch *et al*., 2007, 2008b, *a*, 2010; Koch and Rothwell, 2009; Ziluk *et al*., 2010; Cattaneo and Barchiesi, 2011; Vesia and Davare, 2011; Karabanov *et al*., 2012, 2013; Vesia *et al*., 2013, 2017; Chao *et al*., 2015; Maule *et al*., 2015). The present results cannot be compared to these studies in alert subjects in terms of polarity (inhibition or excitation) nor of task-dependency of the response because of the anaesthesia, but they can be compared in terms of cortical site in the PPC and timing of conditioning stimuli that exert a significant effect onto M1. Our data are compatible with the findings of Karabanov *et al*. (2013) showing two foci along the intraparietal sulcus, the two clusters of active spots along the intraparietal sulcus in the present findings (Figure 3B). The cluster of spots active at 4 ms ISI (Figure 3C) shows striking similarities with the findings of Vesia *et al*. (2013). The most ventral spots along the postcentral sulcus are in the same location as the opercular region that was stimulated in Maule *et al*. (2015). The posterior intraparietal spot stimulated in Koch *et al*. (2007, 2008b) is similar or slightly posterior to the posterior cluster of active spots observed here. Summing up, the current systematic mapping of the PPC is consistent with most of the previous data testing single cortical spots with non-invasive brain stimulation.

### Summary

We show conclusive evidence that in humans direct parieto-motor pathways exist. We investigated motor output to the distal upper limb and found a widespread representation of the hand with seemingly no specific spatial distribution that could parallel the somatotopy of the adjacent somatosensory cortex. Such absence of topographic distribution is well supported by previous data in non-human and human primates that indicate a spatial organization of motor features in the PPC reflecting action types rather than effectors. We did find a specific spatial clustering of motor spots in the PPC according to the polarity of the effect on corticospinal output and to spatial location. A ventral cluster showed excitatory effects, while a dorsal cluster showed inhibitory effects. A third cluster was identified due to its localization, in the mid-portion of the intraparietal sulcus. The clinical neurosurgeon’s attention toward motor function of the parietal lobe is increasingly recognized (Rossi *et al*., 2018). The present data are potentially exploitable as an IONM procedure for the monitoring of complex motor functions, though further investigations are required.

